# Genome report: Genome sequence of the tuliptree scale insect, *Toumeyella liriodendri* (Gmelin)

**DOI:** 10.1101/2024.07.09.602735

**Authors:** Andrew J. Mongue, Amanda Markee, Ethan Grebler, Tracy Liesenfelt, Erin C. Powell

## Abstract

Scale insects are of interest both to basic researchers for their unique reproductive biology and to applied researchers for their pest status. In spite of this interest, there remain few genomic resources for this group of insects. To begin addressing this lack of data, we present the genome sequence of the tuliptree scale insect, *Toumeyella liriodendri* (Gmelin) (Hemiptera: Coccomorpha: Coccidae). The genome assembly spans 536Mb, with over 96% of sequence assembled into one of 17 chromosomal scaffolds. We characterize roughly 66% of this sequence as repetitive and annotate 16,508 protein coding genes. Then we use the reference genome to explore the phylogeny of soft scales (Coccidae) and evolution of karyotype within the family. We find that *T. liriodendri* is an early-diverging soft scale, less closely related to most sequenced soft scales than a species of the family Aclerdidae is. This molecular result bolsters a previous, character-based phylogenetic placement of Aclerdidae within Coccidae. In terms of genome structure, *T. liriodendri* has nearly twice as many chromosomes as the only other soft scale assembled to the chromosome level, *Ericerus pela* (Chavannes). In comparing the two, we find that chromosome number evolution can largely be explained by simple fissions rather than more complex rearrangements. These genomic natural history observations lay a foundation for further exploration of this unique group of insects.

## Introduction

Scale insects are a uniquely interesting group of arthropods, as they hold great importance for both basic and applied research. On the basic side, this group displays remarkable reproductive diversity (Gavrilov 2007; Blackmon *et al*. 2017; Ross *et al*. 2022), from familiar sexual reproduction with chromosomal sex determination (Blackmon *et al*. 2017) to hermaphroditism (Hughes-Schrader 1925; Mongue *et al*. 2021) and a complex form of purely autosomal haplodiploidy known as paternal genome elimination (Nur 1980). Very little is known about either the molecular mechanisms of these alternative sex determination systems, or how the clade transitioned between them. On the applied side, many scale insects are globally invasive pests that require intervention to control (Caltagirone and Doutt 1989; Waterhouse 1991; García Morales *et al*. 2016), but others are both geographically limited and specialized on one or a few host plants (e.g., Spanish moss ensign scale; Morrison 1925). Understanding the factors that differentiate these two groups will help better target screening and management practices. Specifically, both applied and basic research goals are held back by a lack of modern genomic resources compared to other insect clades (e.g., Lepidoptera; Wright *et al*. 2024). In service of beginning to address this scarcity, we have sequenced the genome of the tuliptree scale insect.

*Toumeyella liriodendri* (Gmelin 1790), or tuliptree scale, is a soft scale insect (Hemiptera: Coccomorpha: Coccidae). It reproduces sexually, but with a fully autosomal genome and sex determined by paternal genome elimination (PGE; Nur 1980). Under this system, females are fully diploid in karyotype and gene expression, but males silence and ultimately discard their paternally inherited chromosomes, making them functionally haploid (Gavrilov 2007). Unlike better studied mealybugs (Pseudococcidae; Ross *et al*. 2010; de la Filia *et al*. 2021; Laura Ross *et al*. 2024), which keep the entire silenced paternal genome in somatic cells, soft scales employ a variant of PGE in which some paternal chromosomes are lost during cell division, leading to a variable karyotype between cells within the same male (Gavrilov 2007). Coincident with this rare sex determination system, adults are extremely sexually dimorphic, with large, long-lived, sessile females and small, winged, non-feeding males (Figure 1).

**Figure 1.**
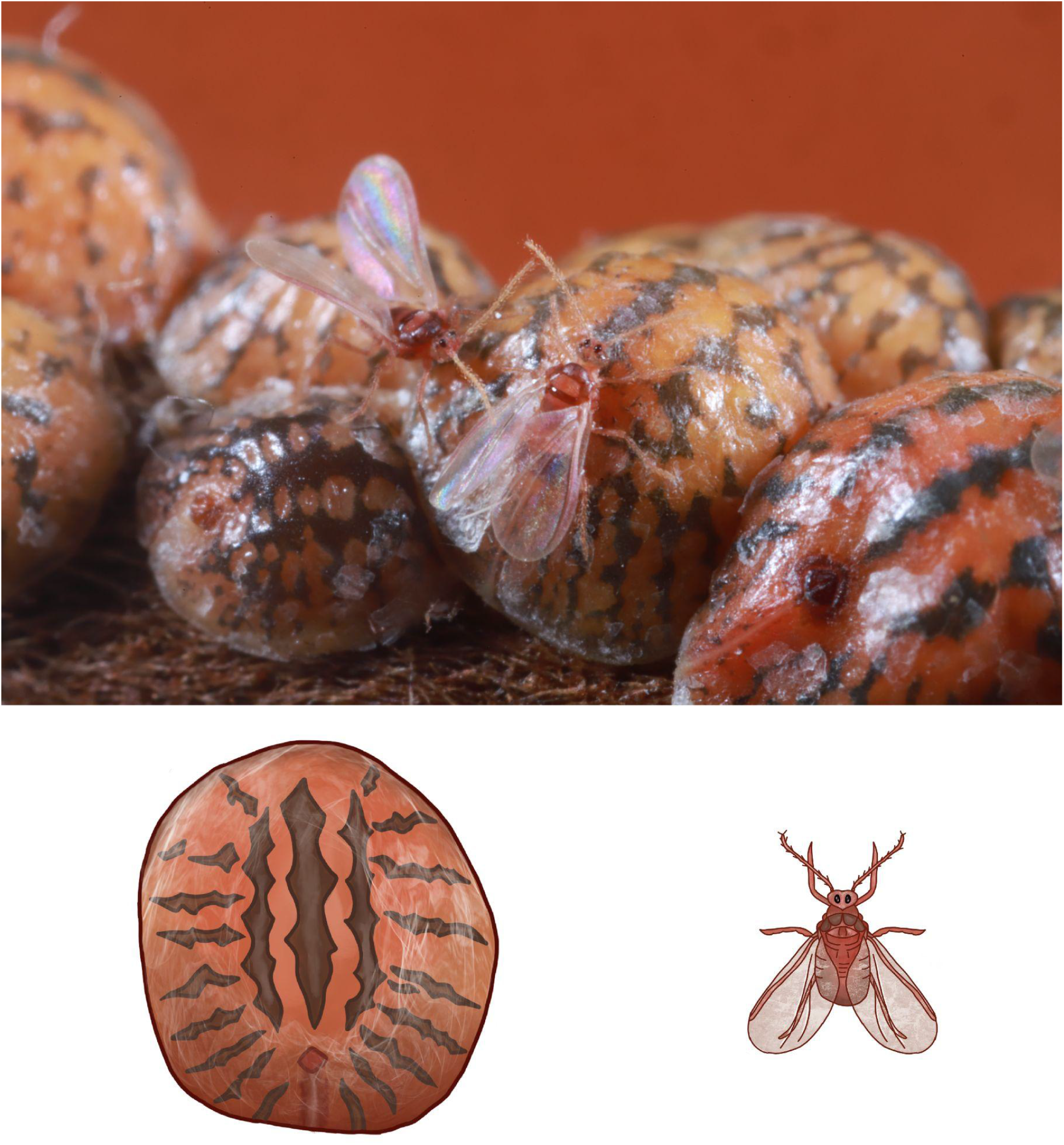
Adult *Toumeyella liriodendri* are incredibly sexually dimorphic. Males (winged) are not only smaller but possess many features (wings, eyes) that are not present in adult females (larger patterned domed individuals). **Top:** Photograph of a colony of *T. liriodendri* with two males and eight females. **Bottom:** Rendering of individual female (left) and male (right) from a dorsal view to highlight sexual dimorphism.

Ecologically, tuliptree scales are presumed native to the eastern United States and likely introduced to California (Hamon and Williams 1984) and Cuba (Novoa *et al*. 2011). This species can be a major pest of native trees including tulip tree, *Liriodendron tulipifera* L., which is commonly grown for timber, and trees in the genus *Magnolia* L., which are popular ornamental and shade trees (Burns and Donley 1970; Hamon and Williams 1984; Gill 1988). While this species prefers Magnoliaceae, it will feed on a variety of hosts across several families (Hamon and Williams 1984; Gill 1988; Miller and Williams 1995).To support study of both the evolution of PGE and monitoring of a tree pest, we present a chromosome-level assembly of *T. liriodendri*.

## Materials and Methods

### Sample collection and identification

We collected adult female *Toumeyella liriodendri* from a single colony feeding on *Magnolia grandiflora* L. (Magnoliaceae) in Gainesville, Florida, United States (29.686122, -82.345128) on June 28^th^, 2023. We confirmed species identity with slide-mounted specimens following Hamon and Williams (1984) and Kondo and Williams (2008). Specimens were deposited in the Florida State Collection of Arthropods (FSCA) in Gainesville, Florida (barcodes FSCA_00072042–FSCA_00072046).

### DNA extraction and sequencing

We followed a sequencing strategy that has proven successful in other recent insect genome sequencing projects, including other scale insects (Laura Ross *et al*. 2024; Mongue *et al*. 2024). First, we extracted DNA from a single adult female using a modified OmniPrep extraction protocol (G-Biosciences, St. Louis, Mo, USA); we followed the manufacturer’s protocol for solid tissue, but extended digestion with proteinase K to overnight (∼15hrs) and extended DNA precipitation step to 1hr at -20 °C. After quality checking, we sequenced genomic DNA at the University of Florida’s Interdisciplinary Center for Biotechnology Research on the PacBio Sequel IIe platform (Menlo Park, Ca, USA) to generate HiFi long reads from a single SMRTCell.

Separately, we pooled adult and juvenile females from the same colony, flash froze them in liquid nitrogen, and sent tissue to Novogene, Inc (Sacramento, Ca, USA) to generate Arima Hi-C linked reads (Carlsbad, Ca, USA) for scaffolding. We completed the initial HiFi sequencing and genome size estimation (below) before HiC sequencing to ensure we chose adequate coverage depth for the latter.

### Genome size estimate

Because little is known about *T. liriodendri* and tissue samples were limited and used for genomic DNA extraction, we sought to estimate the expected genome size directly from the raw PacBio reads by counting k-mer frequencies. To do so, we used Jellyfish v2.3.0 (Marcais and Kingsford 2012) to generate frequencies and then used custom R scripts to estimate genome size from these data.

### Genome assembly

We assembled raw HiFi reads with hifiasm v0.16.1 (Cheng *et al*. 2021), with the “-l 3” stringency parameter for purging duplicated haplotigs. At this stage we performed additional curation steps, both characterizing potential co-bionts and contaminants using blobtools v1.0 (Laetsch and Blaxter 2017) and searching for any remaining haplotigs using the purge_dups tool (https://github.com/dfguan/purge_dups). We also explored circular contigs to identify the mitochondrial sequence and potential co-bionts. We assessed baseline genome completeness with BUSCO v4.1.4 using the hemipteraodb10 dataset (Manni *et al*. 2021). With these steps complete, we aligned Hi-C linked reads to the assembly using Arima’s pipeline (found at https://github.com/ArimaGenomics/mapping_pipeline/): briefly, we independently aligned left and right reads using bwa-mem2 v2.2.1 (Vasimuddin *et al*. 2019) and filtered out chimeric reads while keeping the 5’ end using scripts from the above git repository. We then merged separate alignment files and removed optical duplicates using Picard tools v2.25.5’s MergeSamFiles and MarkDuplicates functions (“Picard toolkit” 2019). We input this curated alignment into YaHS v1.1 (Zhou *et al*. 2023). YaHs generated an initial scaffolded assembly, which we further explored by using the “juicer pre” command to generate JBAT files for visualizing HiC contacts Juicebox 2.17 (Durand *et al*. 2016) for manual correction of the assembly. After visual inspection and correction of misplacements, we saved the updated linkage file and input it into YaHs to run “juicer post” to update the assembly fasta (Zhou *et al*. 2023). We again assessed BUSCO completeness. Based on the failure of the purge_dups approach to identify haplotigs in the primary assembly, we used BUSCO information to screen haplotigs as follows. We concatenated a list of all scaffolds with single copy BUSCO sequences as well as a list of those with multi-copy (duplicate) sequences. We compared the two lists and identified 68 non-chromosomal scaffolds that contained only duplicates and no single-copy BUSCOs. We filtered to remove these from our assembly and proceeded to repeat masking and annotation.

### Repeat masking

We characterized repeats in this final curated genome as follows. First we modeled repeats *de novo* using Repeatmodeler v2.0 (Flynn *et al*. 2020) including a search for long terminal repeats using the “-LTRStruct” parameter. This generated a set of species-specific repeats, which we concatenated to the end of a custom library which consisted of the 2020 Repbase arthropod and hemipteran repeat databases (Bao *et al*. 2015), combined with repeats identified with Repeatmodeler in other high quality genomes: *Icerya purchasi* Maskell (Mongue *et al*. 2024), *Planococcus citri* (Risso) (Laura Ross *et al*. 2024), and another soft scale *Ericerus pela* (Chavannes) (Yang *et al*. 2019). We imported this curated database of hemipteran repeats augmented with scale insect specific and *T. liriodendri* specific repeats into Repeatmasker v4.0.9 (Smit *et al*. 2019) to generate a final soft-masked assembly and summary of repetitive elements.

### Gene Annotation

Lacking enough samples to generate an RNAseq dataset, we chose a de novo approach to gene annotation, using the machine learning tool helixer (Holst *et al*. 2023). Helixer requires only a genome sequence and a general lineage (in our case ‘invertebrate’) to annotate. We passed the softmasked chromosomal assembly to helixer for annotation, but note that the helixer tool claims to not be impacted by presence or absence of masking.

### Phylogeny of Coccidae

We sought to use our newly generated genome to explore phylogenetic relationships between soft scale species. To do so, we downloaded existing datasets for other soft scales and outgroups, as shown in Table 1. The transcriptomic data, we first assembled the transcriptomes using Trinitiy v2.9.0 (Grabherr *et al*. 2011), then ran BUSCO v4.1.4 in transcriptome mode (Manni *et al*. 2021) to extract single copy orthologs for phylogenetic inference; for genome assemblies, we ran BUSCO) directly on the genomes. Next, we used a BUSCO_phylogenomics pipeline (https://github.com/jamiemcg/BUSCO_phylogenomics) to gather single copy BUSCOs present in at least 75% of our sample species and use FastTree v2.1.11 (Price *et al*. 2009) to create individual gene trees for each sequence. Then we used ASTER (https://github.com/chaoszhang/ASTER)’s ASTRAL tool (Zhang *et al*. 2018) to generate a consensus species tree with the “-R” more subsampling and placements options, and choosing *P. citri* to root as the outgroup.

**Table 1.** Summary of currently available soft scale sequence data as well as two outgroups from other families: Aclerdidae and Pseudococcidae. We used this dataset to explore phylogenetic relationships within the family Coccidae.

| Species | Family | Data type | Accession |
| --- | --- | --- | --- |
| *Aclerda* sp. | Acleridae | Transcriptome | SRR1821892 |
| *Planococcus citri* | Pseudococcidae | Genome | GCA_950023065.1 |
| *Ericerus pela* | Coccidae | Genome | GCA_011428145.1 |
| *Ceroplastes cirripediformis* | Coccidae | Transcriptome | SRR1821905 |
| *Coccus* sp. | Coccidae | Transcriptome | SRR1821911 |
| *Parthenolecanium corni* | Coccidae | Genome | GCA_038050395.1 |

### Karyotype Evolution between scale insect families

So far, scale insects of the better-studied mealybug family Pseudococcidae with chromosome-level genome assemblies all have a conserved karyotype of n = 5 (Gavrilov 2007; Li *et al*. 2020; Laura Ross *et al*. 2024), but with additional chromosome-level resources for scale insects comes the opportunity to explore how genome architecture has evolved. We sought to compare our newly generated *T. liriodendri* assembly (n = 17) to that of *E. pela* (n = 9; Chen *et al*. 2021). For this analysis, we removed shorter scaffolds, leaving only the chromosomal pseudomolecule scaffolds and ran Satsuma v2 (Grabherr *et al*. 2010) SatsumaSynteny2 to find orthologous matches across the genome. We then processed these matches with “BlockDisplaySatsuma” and visualized them with “ChromosomePaint” to understand the relationship between karyotypes.

## Results and Discussion

### Sequencing and assembly

We generated 31Gb of raw PacBio HiFi reads (raw data accessions found in Table 2) which we first used to set expectations of genome size using k-mer counting analyses. Our analyses suggested we generated an average of 31x coverage on a roughly 515Mb genome.The karyotype of *T. liriodendri* has not been previously reported, but the contact map of the downstream assembly showed 17 clear linkage groups, even before curation, suggesting the same karyotype as *Neolecanium cornuparvum* (Thro) (Gavrilov 2007). After manual curation of the contact map, 96.5% of the total assembled length (517,894,882 bp) is contained in one of 17 chromosomal scaffolds (Table 3).

**Table 2.** Location of all primary and assembly data generated for *T. liriodendri*.

| Data | Accession |
| --- | --- |
| PacBio HiFi reads | PRJNA1100599 |
| Illumina HiC reads | PRJNA1100599 |
| Assembly | JBEUKQ000000000 |
| Annotation | DOI: 10.5281/zenodo.12611068 |

**Table 3.**
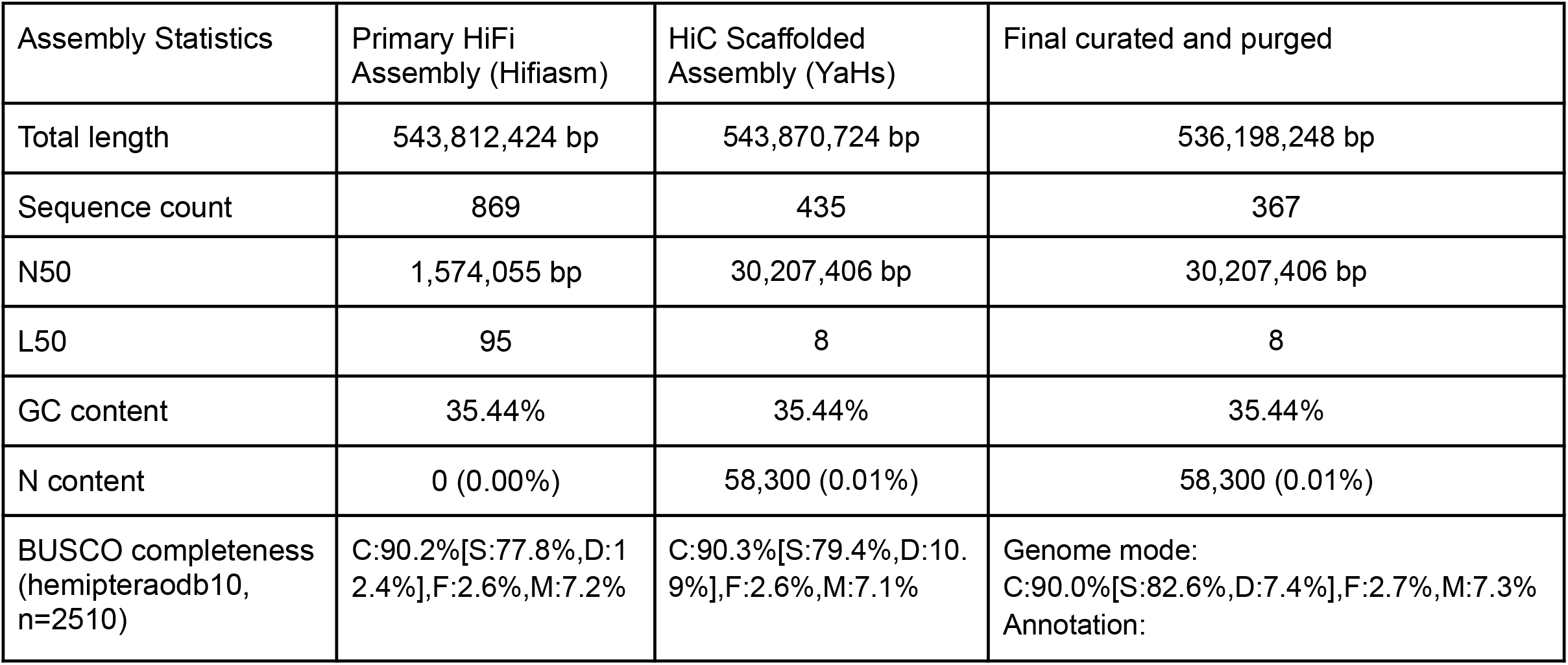
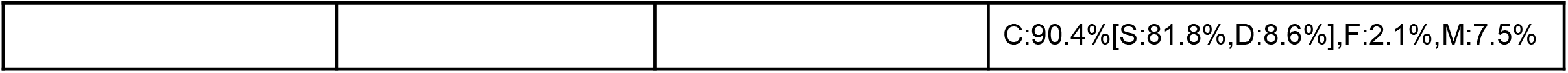
Assembly statistics throughout the assembly process. We report the assembly size, contiguity, and BUSCO completeness for the primary HiFi assembly, the HiC scaffolded assembly, and the final curated assembly reported here.

In the context of other scale insects, there are no closely related species, i.e. congeners, for comparison to set expectations for genome size. That said, the closest well-characterized genome is the soft scale *Ericerus pela*, which has a genome of 650Mb (Yang *et al*. 2019). Other better studied scale insects in the mealybug family Pseudococcidae have genomes of roughly 400Mb (Vea *et al*. 2021; Laura Ross *et al*. 2024). From this perspective the ∼540Mb genome of *T. liriodendri* fits well within the established trend and serves to highlight that the scale insect *Icerya purchasi* is an outlier with a genome of over 1Gb in length (Mongue *et al*. 2024), likely due to its unique reproductive ecology as a self-fertile hermaphrodite (Mongue *et al*. 2021).

Working with the final curated assembly (the far right column of Table 2), we used a combination of *de novo* repeat finding and matching against a database of known hemipteran repeats. In total, we masked 66% of the genome, with results from RepeatMasker summarized in Table 4.This repeat percentage is higher than the ∼55% reported from the related *E. pela* (Yang *et al*. 2019), despite the smaller genome size of *T. liriodendri* but may reflect our use of a larger hemipteran repeat database.

**Table 4.** Summary of masked repeats in the *T. liriodendri* genome. Output is based on the RepeatMasker summary table.

| Repetitive Element | Count | Sequence length (bp) | Percentage of sequence |
| --- | --- | --- | --- |
| SINEs | 3,162 | 529,083 | 0.10 % |
| LINEs | 36,639 | 10,916,041 | 2.04 % |
| LINE1 | 3,326 | 1,083,625 | 0.20 % |
| LINE2 | 1,587 | 515,529 | 0.10 % |
| L3/CR1 | 1,298 | 423,116 | 0.08 % |
| LTR elements | 109745 | 47,806,576 | 8.92 % |
| ERVL | 4 | 253 | 0.00 % |
| ERV Class I | 15834 | 4,356,406 | 0.81 % |
| ERB Class II | 53 | 93,459 | 0.02 % |
| DNA elements | 79,859 | 16,639,012 | 3.10 % |
| hAT-Charlie | 75 | 5,663 | 0.00 % |
| TcMar-Tigger | 13,514 | 1,954,000 | 0.36 % |
| Unclassified | 1,270,206 | 270,951,242 | 50.53 % |
| Total interspersed repeats |  | 34,684,195 | 64.69 % |
| Small RNA | 1,961 | 398,467 | 0.07 % |
| Satellites | 56 | 10,072 | 0.00 % |
| Simple repeats | 86,079 | 4,594,864 | 0.86 % |
| Low complexity | 11,037 | 586,349 | 0.11 % |

Finally, gene content in scale insects is also poorly-characterized. On the low end, annotations of the genome of *E. pela* and *Phenacoccus solenopsis* Tinsley report 12,022 genes (Yang *et al*. 2019) and 11,880 genes (Li *et al*. 2020), respectively. On the high end, species cataloged on the web resource Mealybug Base (https://ensembl.mealybug.org/index.html) range from ∼22,000 (*Pseudococcus longispinus* (Targioni Tozzetti)) to ∼40,000 (*Planococcus citri*) predicted genes. Our annotation of *T. liriodendri* is closer to the low end, with 16,508 protein coding genes. At present, we lack RNA evidence to directly validate our gene predictions, but this lower number is in line with expectations from other insects. Moreover, the 8.6% BUSCO duplication we observe is very similar to the 8.0% observed in the new chromosome-level assembly of *P. citri* (Laura Ross *et al*. 2024), which has yet to be formally annotated.

### Coccidae phylogeny

We used available genomic resources to explore the phylogenetic relationship of sequenced coccid species. Specifically, we used conserved hemipteran BUSCO ortholog sequences to build 2,136 gene trees from which we inferred the overall species tree. We found that *E. pela* is the outgroup to other sequenced soft scales, with *T. liriodendri* being the next branching species (Figure 2). Interestingly, we recover the unspecified *Aclerda* species from Johnson et al. (2018) within the Coccidae, despite its placement in the family Aclerdidae; a larger, mostly morphological phylogeny of scale insects also recovered Aclerdidae within Coccidae (Vea and Grimaldi 2016). This congruence of molecular and morphological results suggest that the classification of Aclerdidae and/or Coccidae may require revision, though it is well beyond the scope of this work.

**Figure 2.**
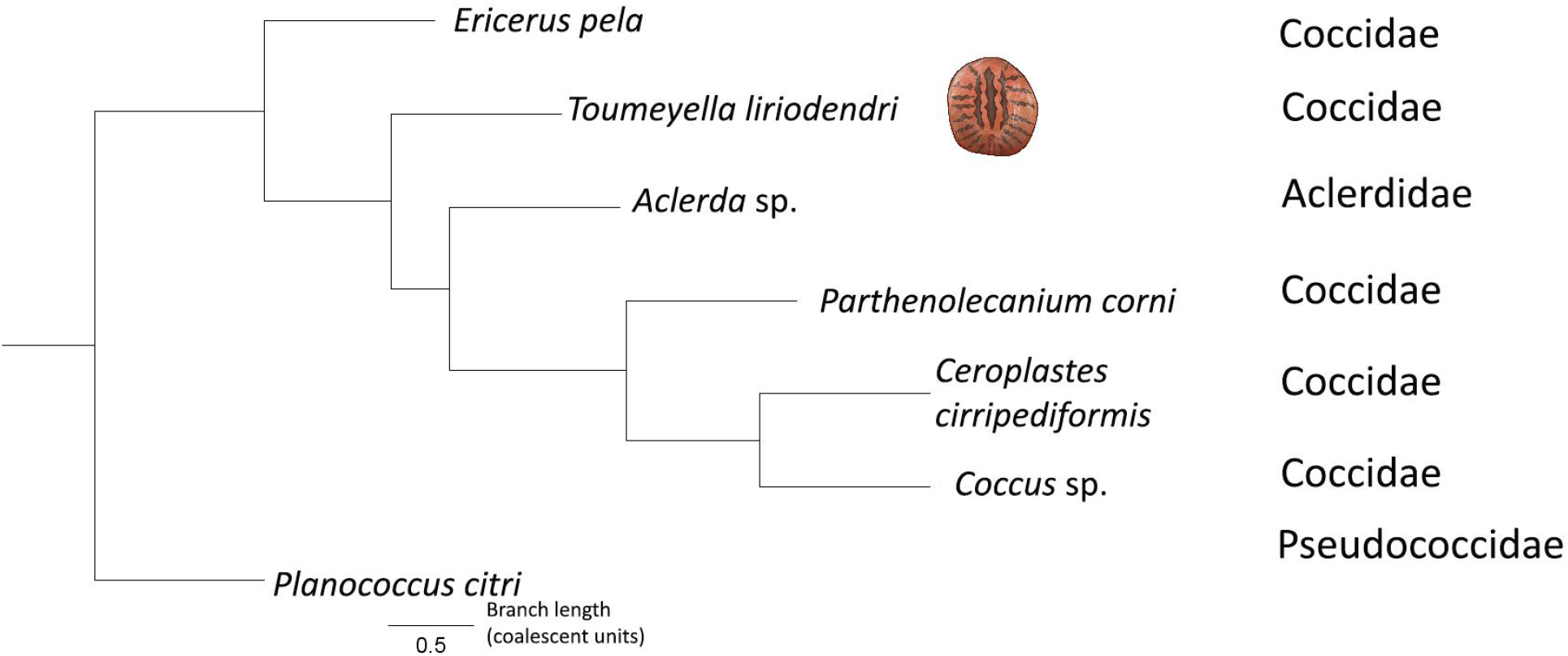
Molecular phylogeny of available Coccidae species. *Toumeyella liriodendri* is an early-diverging soft scale (indicated with a graphic), with *E. pela* as the outgroup to the rest of the family. The only sequenced *Aclerda* species is nested within the family Coccidae. All branches have a local posterior probability of 1.

### Soft scale karyotype evolution

Scale insect genome architecture evolution is poorly studied, but members of Coccidae range in karyotype from n = 5 to n = 18 chromosomes based on currently characterized species (Gavrilov 2007). Based on the karyotypes inferred from genome assemblies (*E. pela* n = 9, *T. liriodendri* n = 17), the simplest explanation for the difference would be a number of fission events in the lineage leading to *T. liriodendri* or fusion events leading to *E. pela*. Our synteny analyses indeed show that most *T. liriodendri* chromosomes are simple fragments of larger *E. pela* chromosomes; in other words the entirety of the *T. liriodendri* chromosome corresponds to a part of a single *E. pela* chromosome. In addition to these simple fissions, there appear to have been at least two fusion events: *T. liriodendri* chromosome 1 and 6 show two distinct blocks that are orthologous to different *E. pela* chromosomes (Figure 3). The question is what drives the evolution of chromosome number between species.

**Figure 3.**
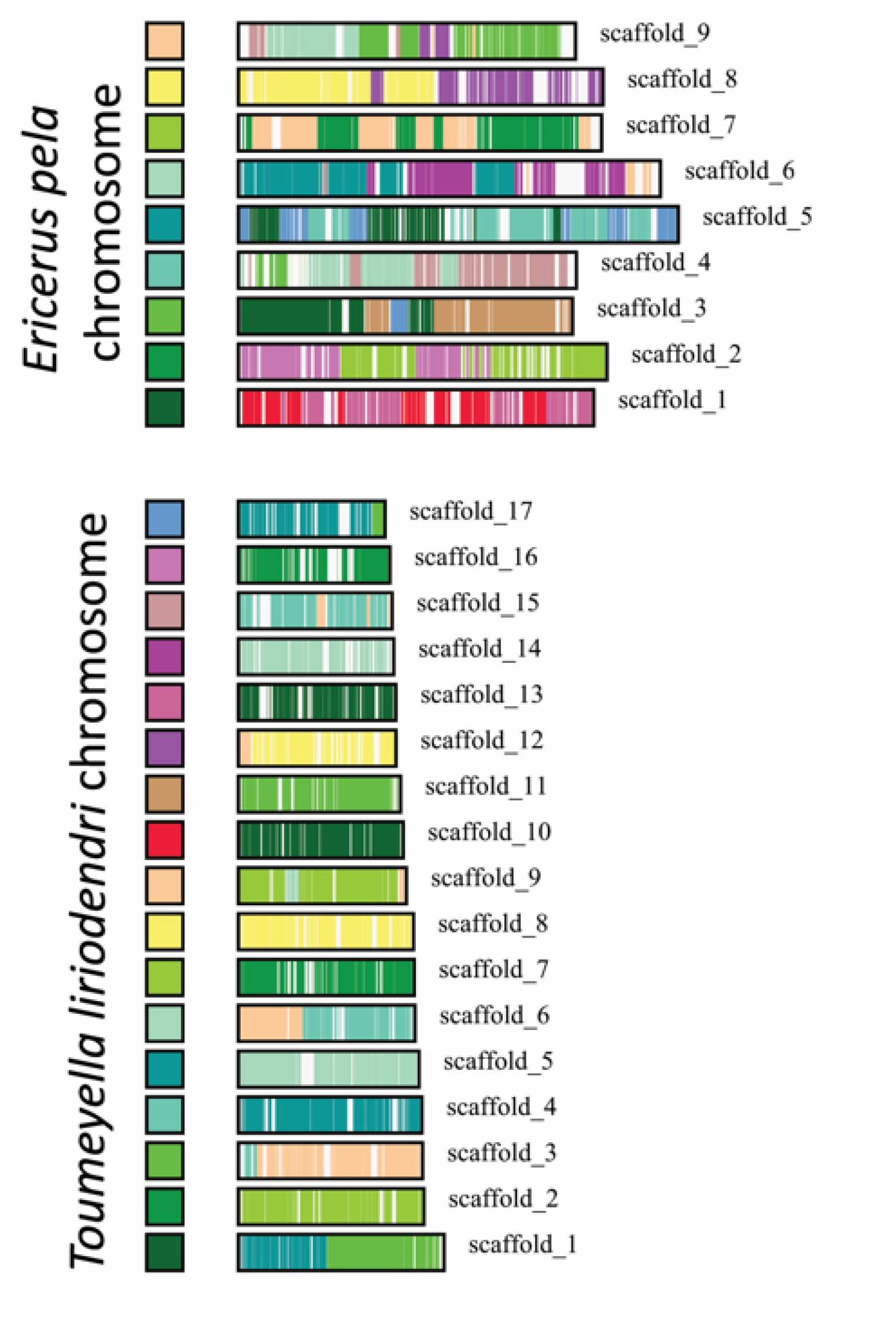
Macrosynteny between the two chromosome-level coccid assemblies. Investigation of synteny between the *E. pela* (n = 9, top) and *T. liriodendri* (n = 17, bottom) genomes. Colored blocks on the left of each chromosome in a given panel encode the color of a syntenic match to a chromosome in the other species (e.g., *T. liriodendri* scaffold 10 (red) matches exclusively to *E. pela* scaffold 1 (dark green)). Overall, karyotype evolution follows a pattern of chromosomal fissions: a given *T. liriodendri* chromosome matches to only one *E. pela* chromosome, but that same *E. pela* chromosome matches to multiple *T. liriodendri* chromosomes.

This variability may be partly explained by the fact that scale insect chromosomes are holocentric, i.e., homologous pairs align along the length of the chromosome rather than at a single centromere point (Parida and Ghosh 1986; Gavrilov 2007). It has been argued that this chromosomal system is more permissive of fusions and fissions, as distinct karyotypes can still successfully align during cell division (Márquez-Corro *et al*. 2019); however, overall rates of chromosome number evolution do not appear to be higher in clades with holocentric taxa (Ruckman *et al*. 2020), and in another group of holocentric insects, the Lepidoptera, chromosome number appears to be remarkably conserved, with the exception of fusions involving the sex chromosome (Mongue *et al*. 2017; Wright *et al*. 2024). Thus other factors must impact the propensity of chromosome number to vary between scale insect species. Indeed, in chromosomally sex-determined taxa, genomic rearrangements involving the sex chromosomes (e.g. X or Z chromosomes) may have some adaptive benefit (Mongue *et al*. 2022) but also come with the cost of potential dosage problems between males and females that requires the evolution of novel gene regulation (Gu *et al*. 2019). In other words, despite a clear adaptive advantage to chromosomal rearrangements, a purely autosomal system such as PGE in Coccidae may be overall more permissive of fusions and fissions, but a broader sampling across scale insect families will be required to test this prediction.

## Conclusions

Here we report a chromosome-level assembly for the ornamental pest tuliptree scale insect, *Toumeyella liriodendri*. This resource will open avenues in both basic research of reproductive diversity of scale insects as well as tracking and management of this ornamental pest. To demonstrate its usefulness, we perform preliminary phylogenetic and synteny analyses with available soft scale data. A deeper exploration of the molecular mechanisms and consequences of paternal genome elimination will require more data, but this genome assembly is a key component.

## Data Availability

All data used in this project are summarized with appropriate accessions and DOIs in Table 1 (for existing data used in comparisons) and Table 2 (newly generated sequence data, genome, annotation data). Ancillary scripts can be found at https://github.com/amongue/ToliGenome/; accessions will become active on publication. For the review process, please use this following link for access to the raw sequence data: https://dataview.ncbi.nlm.nih.gov/object/PRJNA1100599?reviewer=1b4ea566vtao0tsmdnm5k80i ns. The assembly is available via G3 figshare.

## Acknowledgments

Thanks to the University of Florida’s Interdisciplinary Center for Biotechnology Research for support in the PacBio sequencing. Thanks as well to University of Florida Research Computing for providing computational resources and support that enabled the results reported in this publication. URL: http://www.rc.ufl.edu and to the Florida Department of Agriculture and Consumer Services, Division of Plant Industry (FDACS-DPI) for supporting the taxonomy in this work.

